# MPRAnator: a web-based tool for the design of Massively Parallel Reporter Assay experiments

**DOI:** 10.1101/035444

**Authors:** Ilias Georgakopoulos-Soares, Naman Jain, Jesse Gray, Martin Hemberg

## Abstract

DNA regulatory elements contain short motifs where transcription factors (TFs) can bind to modulate gene expression. Although the broad principles of TF regulation are well understood, the rules that dictate how combinatorial TF binding translates into transcriptional activity remain largely unknown. With the rapid advances in DNA synthesis and sequencing technologies and the continuing decline in the associated costs, high-throughput experiments can be performed to investigate the regulatory role of thousands of oligonucleotide sequences simultaneously. Nevertheless, designing high-throughput reporter assay experiments such as Massively Parallel Reporter Assays (MPRAs) and similar methods remains challenging. We introduce MPRAnator, a set of tools that facilitate rapid design of MPRA experiments. With MPRA Motif design, a set of variables provides fine control of how motifs are placed into sequences therefore allowing the user to investigate the rules that govern TF occupancy. MPRA SNP design can be used to investigate the functional effects of single or combinations of SNPs at regulatory sequences. Finally, the Transmutation tool allows for the design of negative controls by permitting scrambling, reversing, complementing or introducing multiple random mutations in the input sequences or motifs.

## Introduction

Temporal and spatial regulation of gene expression is necessary for the functionality of every biological system. Eukaryote cells have many different mechanisms for regulating gene expression, but perhaps the most important one is through transcription factors (TFs). TFs modulate gene expression by binding to their cognate, regulatory motifs. Through the regulation of additional co-factors, the rate of transcription of associated genomic locations is modulated. Experimental methods to determine Protein-DNA interactions, such as Systematic Evolution of Ligands by Exponential Enrichment (SELEX), protein binding microarrays (PBMs) and Chip-Sequencing, have provided valuable information in understanding which genomic sequences a TF has sufficient affinity to bind to (Jolma et al. 2013), (Berger & Bulyk 2009), (Wei et al. 2010), (Chen et al. 2008). However, presence of motif sequence does not necessarily correspond to TF binding and most motif occurrences in the genome are unbound at any given time point (Wang et al. 2012).

Cis-regulatory elements, such as promoters and enhancers, incorporate binding sites for TFs and multi-protein complexes at high density (Zinzen et al. 2009). Typically, multiple TFs will bind to each regulatory element to either activate or repress gene expression in a combinatorial manner. Although computational models have been built based solely on sequence and motif information to predict the location of cis-regulatory elements (Lee et al. 2011) (Yanez-Cuna et al. 2012), disentangling the regulatory information at these loci remains challenging. There does not appear to be a straightforward relationship between the presence of individual TF binding motifs and the level of gene expression of a target gene. Instead, recent studies have implicated multiple genomic features. More specifically, the role of homotypic and heterotypic TF clusters, the contribution of the motif environment, including DNA shape and GC-content and nucleosome positioning have been shown to influence binding (Zinzen et al. 2009), (Dror et al. 2015), (Levo et al. 2015). As a result, there is no simple set of rules that infer TF binding specificity and which determine the regulatory effects of motif occupancy (Slattery et al. 2014).

With increasing amounts of sequencing data available, it has become possible to study the effects of genetic variants on gene expression. In particular, genome-wide association studies (GWAS) have investigated the role of genomic loci on the expression phenotype. One of the most important findings from these studies is that the majority of Single Nucleotide Polymorphisms (SNPs) associated with either an expression phenotype or a disease phenotype are present in non-coding regions of the genome (Maurano et al. 2012). Most of these SNPs are believed to be regulatory, and in some cases the effects of disease-associated non-coding SNPs can be linked to disruption of gene regulation through the disruption of TF binding sites (Spivakov et al. 2012), (Schaub et al. 2012). Nevertheless, determining the functional effect of regulatory SNPs, including investigating the corresponding changes in gene expression levels in a systematic manner remains difficult.

Until recently, *in vitro* systematic examination of regulatory elements has been hindered by the lack of high-throughput assay methods. Fortunately, the rapid decline in oligonucleotide synthesis costs has made the construction of thousands of 100–300bp regulatory sequences for a single microarray affordable. Massively Parallel Reporter Assay (MPRA) experiments use this technology to test how the relative positioning of motifs in reporter assays affect reporter gene expression (Levo & Segal 2014), (Weingarten-Gabbay & Segal 2014), (Melnikov et al. 2012), (Smith et al. 2013), (Dailey 2015), (Inoue & Ahituv 2015), (White et al. 2013). MPRAs are based on microarray synthesis of DNA oligonucleotides, each linked to a unique identifier. The oligonucleotides are then amplified, integrated into plasmids in front of a reporter gene with a minimal promoter and transfected into the cells that one is interested in studying. By measuring the expression levels of the reporter gene using RNA-seq, the regulatory properties of the corresponding sequences can be quantified. Thus, MPRA experiments make it possible to test the regulatory effects of thousands of genomic sequences and their constituent components in a single experiment (Melnikov et al. 2012), (Kheradpour et al. 2013), (White et al. 2013). Even more importantly, MPRAs enable the construction of regulatory modules using synthetic sequences, which are not found in the genome, thereby allowing hypothesis testing about the rules governing gene expression (Sharon et al. 2012), (Mogno et al. 2013), (Smith et al. 2013), (Sharon et al. 2014). For instance, Sharon et al., provided evidence that changes in TF binding site location relative to transcription-start site induce significant changes in gene expression (Sharon et al. 2012) and predicted noise levels of gene expression from constructed regulatory elements (Sharon et al. 2014), while Smith et al. showed that homotypic constructed enhancers confer higher regulatory effects than heterotypic motif clusters (Smith et al. 2013). Moreover, MPRA experiments make it possible to systematically study the regulatory effects of SNPs; thereby relating information provided from GWAS studies at the population level with the exploration of functional effects at the cellular level.

Even though decreasing costs have made MPRA experimental procedure accessible to most labs, widespread adoption of the method is limited by computational challenges. Since each MPRA array can involve tens of thousands of different sequences, it is very hard to manually design MPRA experiments, as there are a plethora of parameters that need to be controlled for. Here we present MPRAnator, a set of tools that allow systematic design of MPRA experiments for the investigation of the effects of SNPs and motifs on regulatory sequences.

**Availability and Implementation:** MPRAnator tools are implemented in Python and Javascript and are freely available at: www.genomegeek.com. The source code is available on www.github.com/hemberg-lab/MPRAnator/ under the MIT license.

## 2. Features and Description

The MPRAnator tool set is implemented in Python using the Django framework and javascript for the web interface. The REST API allows programmatic access to MPRAnator using simple URLs.

MPRAnator allows users to systematically design synthetic DNA sequences for high throughput experiments in an interactive manner. Currently, MPRAnator provides support for three different types of investigations regarding the effect of regulatory sequences. The MPRA Motif design tool can be used to systematically generate synthetic sequences with single motifs or combinations of motifs placed at preselected positions. The MPRA SNP design tool can be used to investigate the regulatory effects of single or combinations of SNPs. The Transmutation tool allows for the design of different types of negative controls for MPRA experiments.

### 2.1. MPRAnator Motif Design Tool

The MPRAnator Motif Design Tool requires a minimum of two inputs: i) a set of FASTA scaffold sequences and ii) a list of motifs in FASTA format. It outputs a set of FASTA sequences, where a subset of the nucleotides in the scaffold was substituted for the different motifs that were provided by the user. A set of optional inputs provides the user with fine control of how the motifs are placed. The user can adjust the frequency of motif substitutions into the sequences and use restriction parameters to control the relative distance between motifs substituted in the sequence as well as the range within the sequence at which it is permitted to substitute motifs (Fig. 1). Moreover, the user can select to incorporate uniquely identifiable barcodes to the output sequences, select the barcode size (set to zero to exclude barcodes), select the minimum Levenshtein distance between barcodes and restrict the range of the barcode GC content. Furthermore, the user can select the multiplicity of each sequence to optimize the experiment. On the website all parameters are set to sensible default values. To facilitate integration of the sequence into a vector, restriction sites, adaptor sequences or other sequences of interest can be added. In the output file, the headers of the scaffold sequences contain the corresponding information including the motif sequences, the positions of substitution and any additional subcomponents added to the final product. If any of the restriction sites are identified in any generated oligonucleotide sequence they are reported in the header of the corresponding sequence. To facilitate visualization, the substituted motifs are colour-marked. Importantly, the tool supports a modular design and the order of the constituent parts can be easily altered using a drag and drop interface (Fig. 3). MPRAnator allows for a very high degree of flexibility in the design of the experiment and it is straightforward for the user to explore a wide range of sequence layouts. The flexible layout means that the output sequences are not restricted to MPRA experiments, but they can easily be linked to similar protocols such as BunDLE-seq (Levo et al. 2015).

**Figure1:**
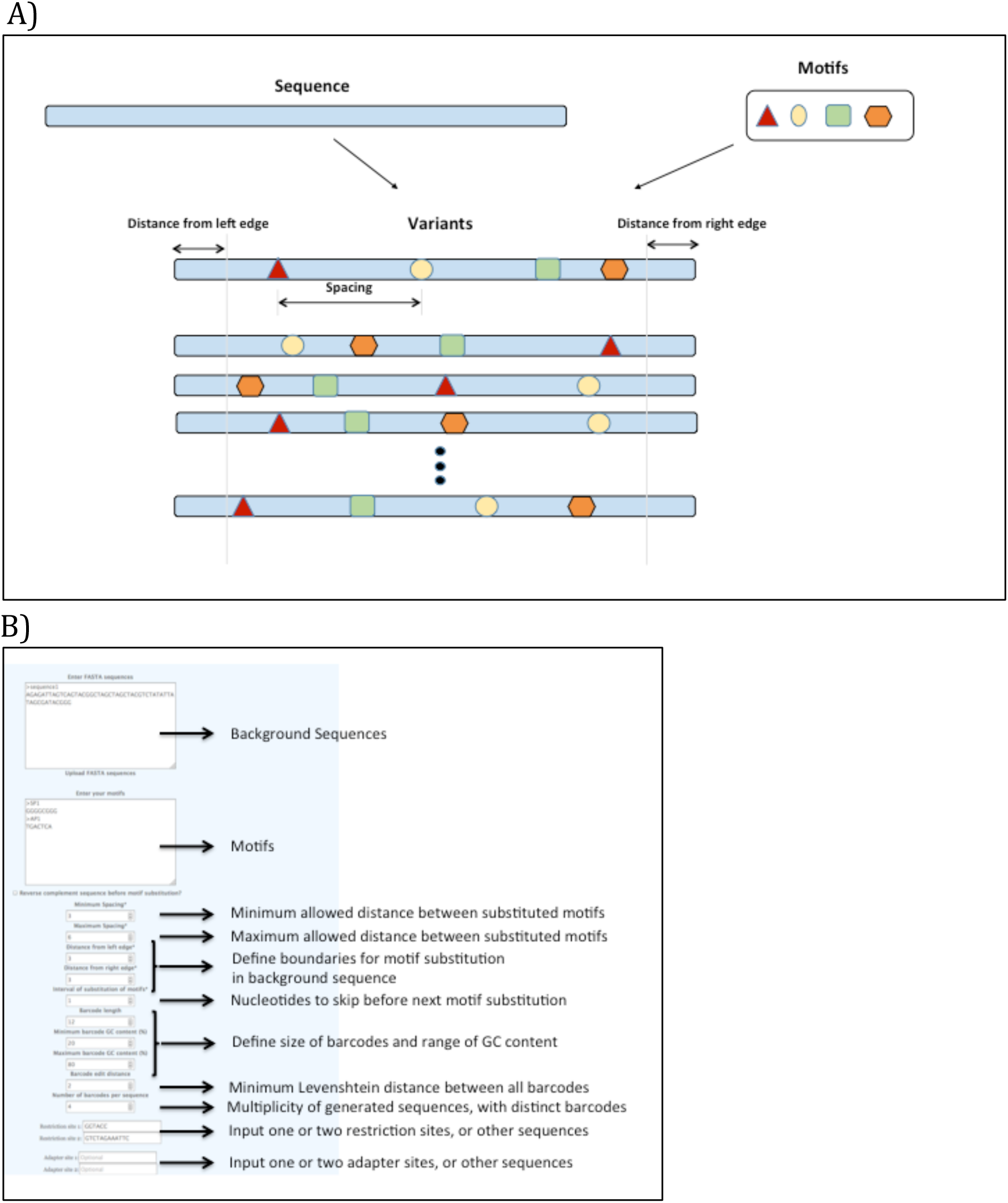
A) MPRAnator Motif design. Up to 4 motifs are substituted into each of the background sequences using all possible permutations. Distance from the edges and minimum and maximum spacing between motifs restrict the positioning of motifs in the sequences. Interval of substitution (not shown) determines the frequency of motif substitutions in consecutive sequences (substituted every X nucleotides within the inputted sequence). B) Screenshot from MPRAnator Motif design query page, with explanation of each variable.

### 2.2 MPRAnator SNP design tool

The MPRAnator SNP design tool requires a minimum of two inputs: i) a set of FASTA scaffold sequences and ii) a list of associated SNPs represented using the variant call file (VCF) format. The VCF format supports the use of up to 12 columns for each locus, but MPRAnator only uses the information found in the first 5 columns. For each sequence in the FASTA file, MPRAnator will substitute the associated SNPs. If more than one SNP is found at a given locus, all combinations will be generated (Fig. 2). Deletions or insertions (up to 10nt) in VCF format are also allowed. Since several methods of oligonucleotide synthesis work best when all oligos have similar lengths, the instances with an insertion must be trimmed while the sequence with a deletion must be expanded. MPRAnator solves this problem by adding adenines to one end of the sequences that are too short, and by trimming one end of the sequences that are too long (Fig. 2).

**Figure2:**
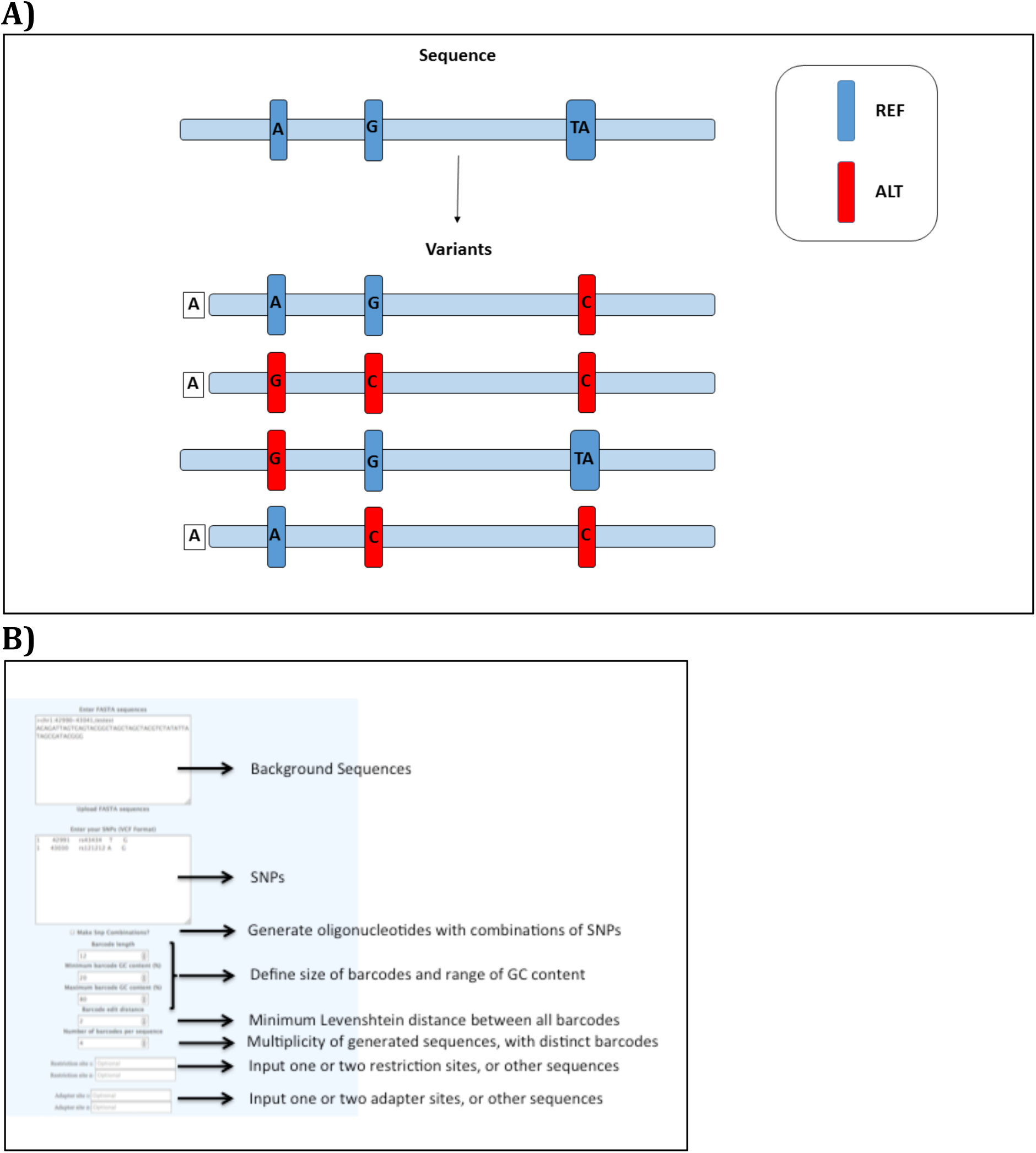
A) MPRAnator SNP design. For inserted sequences, variants are generated that contain the associated SNPs. If multiple SNPs are found in a sequence the user can select to also generate oligonucleotides with their combinations. For deletions, adenines are added at the edge of the sequence and for insertions the sequence is trimmed to maintain the same length across all output sequences. B) Screenshot from: MPRAnator SNP design query pages with explanation of each variable.

As with the motif design tool, a set of optional inputs can be selected for optimal design of the experiment. The user can select to incorporate uniquely identifiable barcodes to the output sequences, select the barcode size (set to zero to exclude barcodes), minimum Levenshtein distance between barcodes and restrict the range of the barcode GC content. Although the default is to substitute all combinations of SNPs in the input sequences, there is also the option to substitute only a single SNP at a time. To facilitate integration of the sequence to a vector, restriction sites, adaptor sequences or other sequences of interest can be added. The headers of the scaffold sequences contain all the corresponding information, including the SNP names, their position and their sequence as well as any additional subcomponents added into the final product. If any of the restriction sites that have been added to the scaffold sequences are identified in any generated oligonucleotide sequence they are reported in the header of the corresponding sequence. Finally, the order of the constituent parts can be altered using a drag and drop interface, allowing flexibility in the design of the experiment (Fig. 3).

**Figure3:**
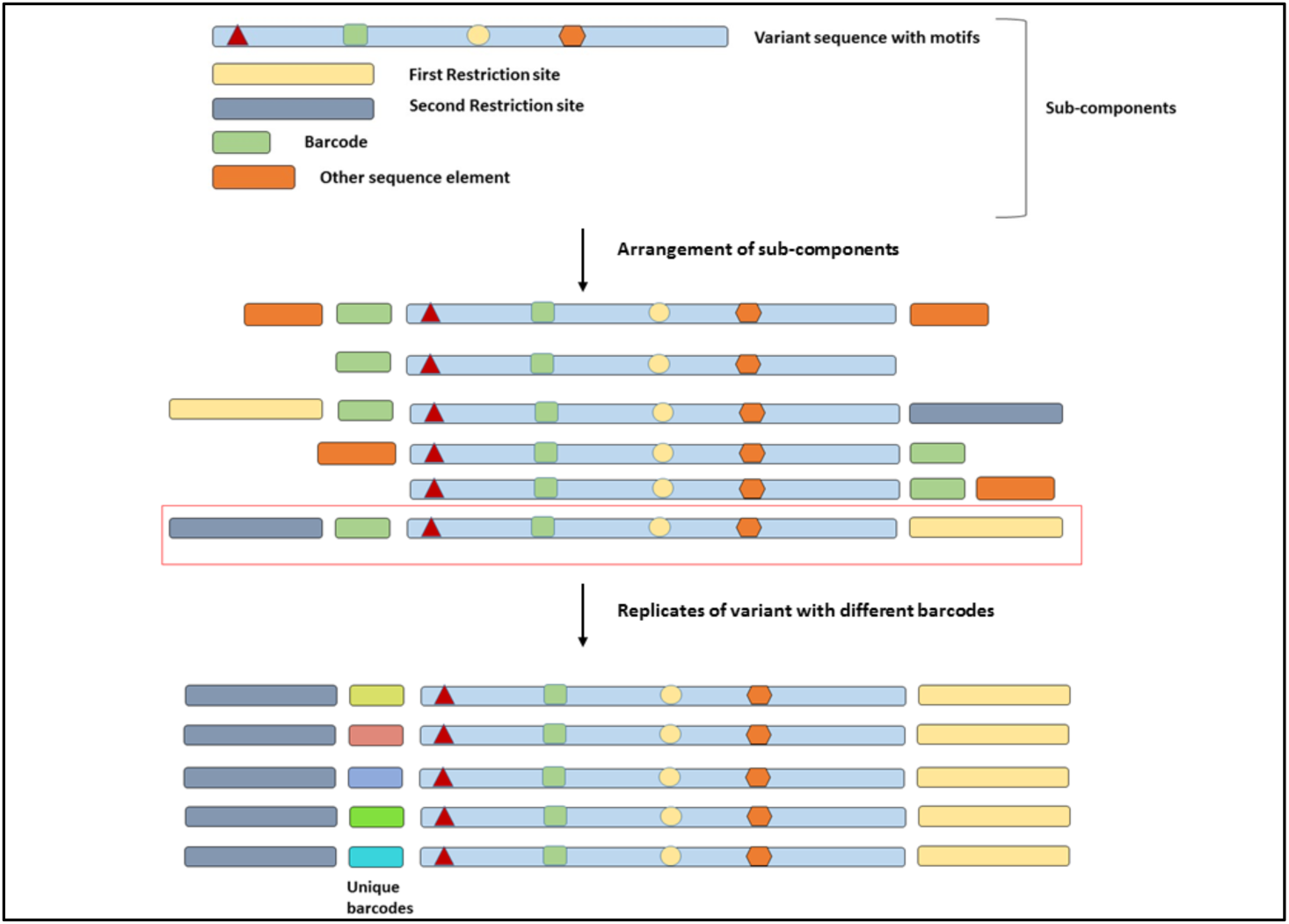
Modular design for final output. Sub-components can be placed in any order. Replicates of each sequence can be generated, each with distinct barcode sequence.

Screenshots of MPRAnator SNP and Motif design webpages are annotated to explain all variables that can be controlled during the design of MPRA experiments (Figures 1B, 2B).

### 2.3 Transmutation tool

The transmutation tool can be used to modify a set of sequences or motifs. The software includes four options: i) scramble, ii) reverse, iii) complement a set of motifs or sequences or iv) introduce multiple random mutations to destroy a motif or sequence s functionality, therefore serving to generate negative controls for the MPRA SNP and Motif design tools. The transmutation tool takes a FASTA file as an input and it will output another FASTA file. The output file will either have a specific number of point mutations per sequence, or each sequence will have been scrambled through a random permutation of the nucleotides or reversed or complemented. Information regarding mutations, reversing, complementing and scrambling for each sequence is stored in the headers of the output FASTA file. Mutated nucleotides are colour-marked for visualization purposes.

## Summary

MPRA is a new and exciting technology, which makes it possible to study how gene expression is regulated in a high-throughput manner. We have presented MPRAnator, the first MPRA design tool. MPRAnator provides an intuitive, interactive, web-based user-interface, which allows the user to systematically construct experiments to study the effects of motifs and SNPs. The regulatory effects of both motifs and SNPs can be studied in isolation as well as combinatorially. The MPRAnator tool set is highly flexible allowing for the incorporation of other genomic sequences as sub-components, the selection of ordering of the sub-components into the final experimental design. The generated sequences can be incorporated into different types of vectors, such as viruses or plasmids, and we believe that MPRAnator can be used for other types of high-throughput designs besides MPRA experiments. The user-friendly nature of MPRAnator will facilitate the adoption of the MPRA technology and allow the method to be used more widely within the scientific community.

## REFERENCES

Berger, M.F. & Bulyk, M.L., 2009. Universal protein-binding microarrays for the comprehensive characterization of the DNA-binding specificities of transcription factors. Nature protocols, 4, pp.393–411. Available at: http://dx.doi.org/10.1038/nprot.2008.195.

Chen, C.Y. et al., 2008. Discovering gapped binding sites of yeast transcription factors. Proceedings of the National Academy of Sciences, 105, p.2527.

Dailey, L., 2015. High throughput technologies for the functional discovery of mammalian enhancers: New approaches for understanding transcriptional regulatory network dynamics. Genomics. Available at: http://www.ncbi.nlm.nih.gov/pubmed/26072436.

Dror, I. et al., 2015. A widespread role of the motif environment on transcription factor binding across diverse protein families. Genome Res. Available at: http://dx.doi.org/10.1101/gr.184671.114.

Inoue, F. & Ahituv, N., 2015. Decoding enhancers using massively parallel reporter assays. Genomics. Available at: http://linkinghub.elsevier.com/retrieve/pii/S0888754315300082.

Jolma, A. et al., 2013. DNA-binding specificities of human transcription factors. Cell, 152, pp.327–339. Available at: http://dx.doi.org/10.1016/j.cell.2012.12.009.

Kheradpour, P. et al., 2013. Systematic dissection of regulatory motifs in 2000 predicted human enhancers using a massively parallel reporter assay. Genome research, 23, pp.800–811. Available at: http://www.pubmedcentral.nih.gov/articlerender.fcgi?artid=3638136&tool=pmcentrez&rendertype=abstract.

Lee, A.P., Brenner, S. & Venkatesh, B., 2011. Mouse transgenesis identifies conserved functional enhancers and cis-regulatory motif in the vertebrate LIM homeobox gene Lhx2 locus. PLoS ONE, 6, p.e20088. Available at: http://www.pubmedcentral.nih.gov/articlerender.fcgi?artid=3100342&tool=pmcentrez&rendertype=abstract.

Levo, M. et al., 2015. Unraveling determinants of transcription factor binding outside the core binding site. Genome Research, p.gr.185033.114. Available at: http://genome.cshlp.org/content/early/2015/03/11/gr.185033.114?top=1.

Levo, M. & Segal, E., 2014. In pursuit of design principles of regulatory sequences. Nature reviews. Genetics, 15, pp.453–68. Available at: http://www.ncbi.nlm.nih.gov/pubmed/24913666.

Maurano, M.T. et al., 2012. Systematic Localization of Common Disease-Associated Variation in Regulatory DNA. Science, 337, pp.1190–1195. Available at: http://www.pubmedcentral.nih.gov/articlerender.fcgi?artid=3771521&tool=pmcentrez&rendertype=abstract.

Melnikov, A. et al., 2012. Systematic dissection and optimization of inducible enhancers in human cells using a massively parallel reporter assay. Nature biotechnology, 30, pp.271–7. Available at: http://www.nature.com/nbt/journal/v30/n3/full/nbt.2137.html?WT.ec_id=NBT-201203.

Mogno, I., Kwasnieski, J.J.C.J.J.C. & Cohen, B. a., 2013. Massively parallel synthetic promoter assays reveal the in vivo effects of binding site variants. Genome research, 23(11), pp.1908–15. Available at: http://www.ncbi.nlm.nih.gov/pubmed/23921661\nhttp://genome.cshlp.org/content/early/2013/08/06/gr.157891.113.abstract.

Schaub, M.A. et al., 2012. Linking disease associations with regulatory information in the human genome. Genome research, 22, pp.1748–1759. Available at: http://genome.cshlp.org/content/22/9Z1748.full.

Sharon, E. et al., 2012. Inferring gene regulatory logic from high-throughput measurements of thousands of systematically designed promoters. Nature biotechnology, 30(6), pp.521–30. Available at: http://www.ncbi.nlm.nih.gov/pubmed/22609971.

Sharon, E. et al., 2014. Probing the effect of promoters on noise in gene expression using thousands of designed sequences. Genome Research, p.gr.168773.113-. Available at: http://genome.cshlp.org.gate1.inist.fr/content/early/2014/07/16/gr.168773.113.abstract.

Slattery, M. et al., 2014. Absence of a simple code: how transcription factors read the genome. Trends in Biochemical Sciences, 39, pp.381–399. Available at: http://www.sciencedirect.com/science/article/pii/S0968000414001212.

Smith, R.P. et al., 2013. Massively parallel decoding of mammalian regulatory sequences supports a flexible organizational model. Nat Genet, 45, pp.1021–1028. Available at: http://www.ncbi.nlm.nih.gov/pubmed/23892608.

Spivakov, M. et al., 2012. Analysis of variation at transcription factor binding sites in Drosophila and humans. Genome biology, 13, p.R49. Available at: http://genomebiology.com/2012/13/9/R49.

Wang, J. et al., 2012. Sequence features and chromatin structure around the genomic regions bound by 119 human transcription factors. Genome research, 22, pp.1798–1812. Available at: http://www.pubmedcentral.nih.gov/articlerender.fcgi?artid=3431495&tool=pmcentrez&rendertype=abstract.

Wei, G.-H. et al., 2010. Genome-wide analysis of ETS-family DNA-binding in vitro and in vivo. The EMBO journal, 29, pp.2147–60. Available at: http://www.pubmedcentral.nih.gov/articlerender.fcgi?artid=2905244&tool=pmcentrez&rendertype=abstract.

Weingarten-Gabbay, S. & Segal, E., 2014. The grammar of transcriptional regulation. Human Genetics, 133, pp.701–711.

White, M. a et al., 2013. Massively parallel in vivo enhancer assay reveals that highly local features determine the cis-regulatory function of ChIP-seq peaks. Proceedings of the National Academy of Sciences of the United States of America, 110, pp.11952–7. Available at: http://www.pubmedcentral.nih.gov/articlerender.fcgi?artid=3718143&tool=pmcentrez&rendertype=abstract.

Yanez-Cuna, J.O. et al., 2012. Uncovering cis-regulatory sequence requirements for context specific transcription factor binding. Genome research, 22, pp.2018–2030. Available at: http://genome.cshlp.org/cgi/doi/10.1101/gr.132811.111\npapers3://publication/doi/10.1101/gr.132811.111.

Zinzen, R.P. et al., 2009. Combinatorial binding predicts spatio-temporal cis-regulatory activity. Nature, 462, pp.65–70. Available at: http://dx.doi.org/10.1038/nature08531.

